# Early enrichment of adaptive NK cells during acute HIV-1 infection is associated with subsequent durable viral control

**DOI:** 10.1101/2025.03.17.643703

**Authors:** Aljawharah Alrubayyi, Amin S. Hassan, Anthony Hsieh, Jonathan Hare, Jill Gilmour, Matt A Price, William Kilembe, Etienne Karita, Eugene Ruzagira, Joakim Esbjörnsson, Eduard J. Sanders, Sarah L. Rowland-Jones, Dimitra Peppa

## Abstract

Early immune events during acute HIV-1 infection (AHI) can shape subsequent viral control, yet the innate immune responses preceding durable control remain poorly defined in humans. Here, we leveraged a historical longitudinal cohort recruited near peak HIV-1 viraemia and followed for up to six years prior to initiation of antiretroviral therapy (ART). We found an early enrichment of FcεRIγ CD57 adaptive NK cells during AHI, accompanied by increased granzyme B expression and lower Ki-67 expression than in canonical NK cells. Mechanistically, *in vitro* modelling showed that inflammatory cytokines, in the presence of plasma collected from participants with AHI and HIV-1-infected targets, induced FcεRIγ NK cell expansion, consistent with an antibody-dependent mechanism. Adaptive NK cells degranulated more strongly than canonical NK cells against autologous CD4 T cells infected with transmitted/founder virus. Furthermore, we showed that higher frequencies of adaptive NK cells during the first month of infection distinguished individuals who subsequently maintained plasma VL <10,000 copies ml ¹ throughout follow-up. Together, these data highlight adaptive NK cells as a candidate early immune correlate of durable HIV control and a population to target in vaccine strategies that engage Fc receptors.

## Main

Durable control of HIV-1 in the absence of antiretroviral therapy (ART) remains a central goal for HIV cure and vaccine strategies. The first weeks after infection are a critical period in which viraemia rises rapidly and antiviral immune responses emerge; yet it remains unclear which early cellular responses predict long-term control. Further insight into the multifaceted virus-host interactions during acute HIV-1 infection (AHI) could help identify future prophylactic and therapeutic targets but is difficult to address in contemporary cohorts because ART is now recommended for prompt initiation after HIV diagnosis. Historical cohorts, recruited before universal immediate ART, with samples collected near peak viraemia and prolonged untreated follow-up, offer a rare opportunity to investigate how early immune events shape subsequent viral outcomes in natural HIV-1 infection.

Natural killer (NK) cells are among the earliest immune populations to respond during HIV-1 infection and can exert antiviral activity through both cytokine-mediated and antibody-dependent mechanisms ^1,2^. Among NK cell subsets, terminally differentiated adaptive NK cells are of particular interest. In Human cytomegalovirus (HCMV) infection, they expand in response to inflammatory cytokines and viral stimuli and display enhanced antibody-dependent effector function ^3–5^. Their expansion requires IL-12 ^3^ and can be driven by CD16 engagement of antibody-coated targets ^5–7^. These adaptive NK cells, which are typically characterised by NKG2C expression and loss of FcεRIγ, are enriched in people living with HIV-1 (PLWH) ^8–10^, and have been associated with spontaneous HIV control ^11–14^. More recently, higher frequencies of NKG2C adaptive NK cells have been associated with lower frequencies of intact proviruses during long-term suppressive ART ^15^. However, the dynamics of adaptive NK cells during the first weeks of infection and their role in subsequent viral control remain poorly defined.

Here, we drew on the International AIDS Vaccine Initiative (IAVI) Protocol C cohort, recruited between 2006 and 2011, in which participants were enrolled as early as two weeks after the estimated date of infection (EDI) and followed without ART for up to six years ^16^. Using longitudinal immunophenotyping, plasma cytokine profiling, an *in vitro* expansion assay and autologous HIV-1-infected target-cell models, we examined early NK cell responses during acute HIV-1 infection, identified signals associated with NK cell expansion, and assessed whether the magnitude of the early adaptive NK cell response was linked to subsequent viral control.

### Longitudinal profiling of a historical acute HIV-1 infection cohort

We studied 50 participants from the IAVI Protocol C cohort recruited at four research centres in Kenya, Rwanda, Uganda and Zambia (**Fig. 1a and Methods**). Participants were selected based on the availability of PBMC samples at both one and three months post estimated date of infection (EDI) and were followed up to six years without ART according to local treatment guidelines at the time of recruitment ^16,17^. The cohort included participants with different HIV-1 subtypes (n=27 subtype A, n=14 subtype C, and n=9 subtype D; **Fig. 1a and Supplementary Table 1**). Longitudinal analyses used samples collected at two weeks, one month, and three months post-EDI, with samples available from 12 participants at two weeks and all 50 participants at one and three months. Viral load and CD4 T-cell counts were available for up to six years post-EDI without ART, enabling assessment of early immune responses in relation to subsequent untreated viral trajectories (**Fig. 1a-c**). The number of participants contributing ART-naive measurements declined over follow-up to 32 at 2.5 years, 19 at 3.5 years and 5 at 6 years, reflecting ART initiation and loss to follow-up (**Supplementary Table 2**). All measurements after ART initiation were excluded. Consistent with previous AHI reports ^18^, plasma viral load rose rapidly to peak levels during the first weeks post-EDI and then declined toward viral set point (**Fig. 1b**). CD4 T-cell counts were broadly stable across the acute phase and declined modestly over the subsequent two years (**Fig. 1c**).

**Figure 1.**
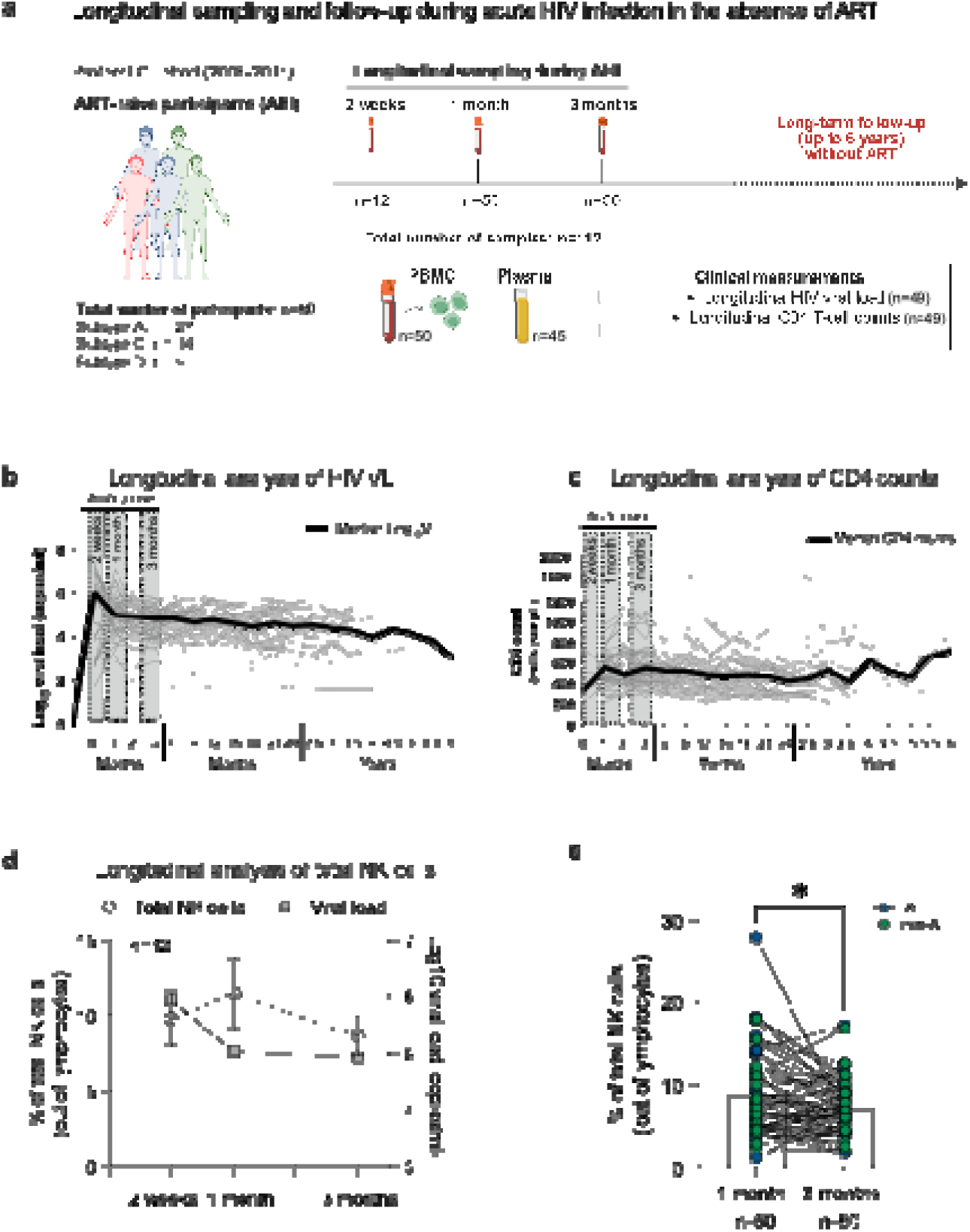
Longitudinal sampling and clinical follow-up of a historical ART-naïve acute HIV-1 infection cohort. **(a)** Study design of the IAVI Protocol C cohort. Longitudinal study design, timepoints, and number of PBMC samples collected at each timepoint (total number of participants n=50). **(b)** Longitudinal analysis of plasma VL for participants following the onset of detectable plasma viraemia and up to six years. All measurements after the initiation of ART were excluded from the analysis. The thickened line represents the median log_10_ viral load. The timepoints sampled in this study are highlighted in the grey-shaded box (1^st^ timepoint = two weeks; 2^nd^ timepoint = one month; 3^rd^ timepoint = three months). **(c)** Longitudinal analysis of CD4^+^ T-cell counts. **(d)** Longitudinal analysis of total NK cells (dashed black line) and log_10_ viral loads (grey solid line) from two weeks and up to three months post-infection in a subgroup of participants (*n*=12). **(e)** Summary analysis of total NK cells in all participants. Blue dots denote subtype A participants, and green dots denote non-A subtype participants (A: *n*=27 and non-A: *n* =23). The number of participants included in longitudinal VL analysis (**b and c**) is reported in Supplementary Table 2. Significance was determined by two-tailed Wilcoxon matched-paired signed rank test. *p<0.05.

Given previous reports of more favourable clinical outcomes in HIV-1 subtype A infection in this cohort, ^19,20^ we examined clinical trajectories by infecting subtype (**Extended Data Fig. 1**). Participants with subtype A and non-A infection showed broadly similar early VL patterns, with an initial peak followed by a decline, but their later VL trajectories diverged (**Extended Data Fig. 1a-c**). VL was lower in the subtype A participants at 18 months and 2.5 years post-EDI (**Extended Data Fig. 1d**), with no significant differences at the other tested timepoints (**Supplementary Table 2**). CD4 T-cell trajectories showed considerable variation within both groups, with subtype A participants having higher CD4 T-cell counts at 2.5 years (**Extended Data Fig. 1e,f and Supplementary Table 2**). NK cell responses were compared across the full cohort, with infecting subtype shown in the figures.

### Progressive enrichment of adaptive NK cells during acute HIV-1 infection

We first performed longitudinal NK cell analysis to define the evolution of NK cell responses during untreated acute HIV-1 infection. Total NK cell proportions were followed alongside plasma viraemia in the subgroup sampled at all three timepoints (**Fig. 1d**) and decreased significantly from one to three months in paired samples from the full cohort of 50 participants, despite variation between individuals (**Fig. 1e**). There were no significant differences in the frequencies of the main NK cell subsets (**Extended Data Fig. 2a-c**).

To further characterise NK cell responses during AHI, we performed high-dimensional analysis of the more abundant CD56^dim^ NK cells using the maturation (CD57, NKG2A, KIRs), activation (CD38 and CD2), and adaptive (NKG2C, PLZF, Siglec-7, and FcεRIγ) markers (**Extended Data Fig. 3a-c**). This analysis showed a highly differentiated population with an adaptive phenotype, NKG2C CD2 PLZF KIR Siglec-7 FcεRIγ CD57 (metacluster 5), which comprised up to 46% of the CD56^dim^ compartment and increased in abundance over time (**Extended Data Fig. 3a-c**). Consistent with this, manual gating showed that FcεRIγ CD57 adaptive NK cells were enriched over the course of AHI, with frequencies increasing significantly from one to three months post-EDI (**Fig. 2a,b**). Further analysis showed that NKG2C CD56^dim^ NK cells were predominantly FcεRIγ, with FcεRIγ NKG2C cells significantly outnumbering FcεRIγ NKG2C cells at all three timepoints **(Extended Data Fig. 3d)**. This is in keeping with previous studies showing that human adaptive NK cell populations are commonly characterised by FcεRIγ loss and enrichment of NKG2C expression ^5,21^, and consistent with the co-expression of these markers within metacluster 5 (**Extended Data Fig. 3c**).

**Figure 2.**
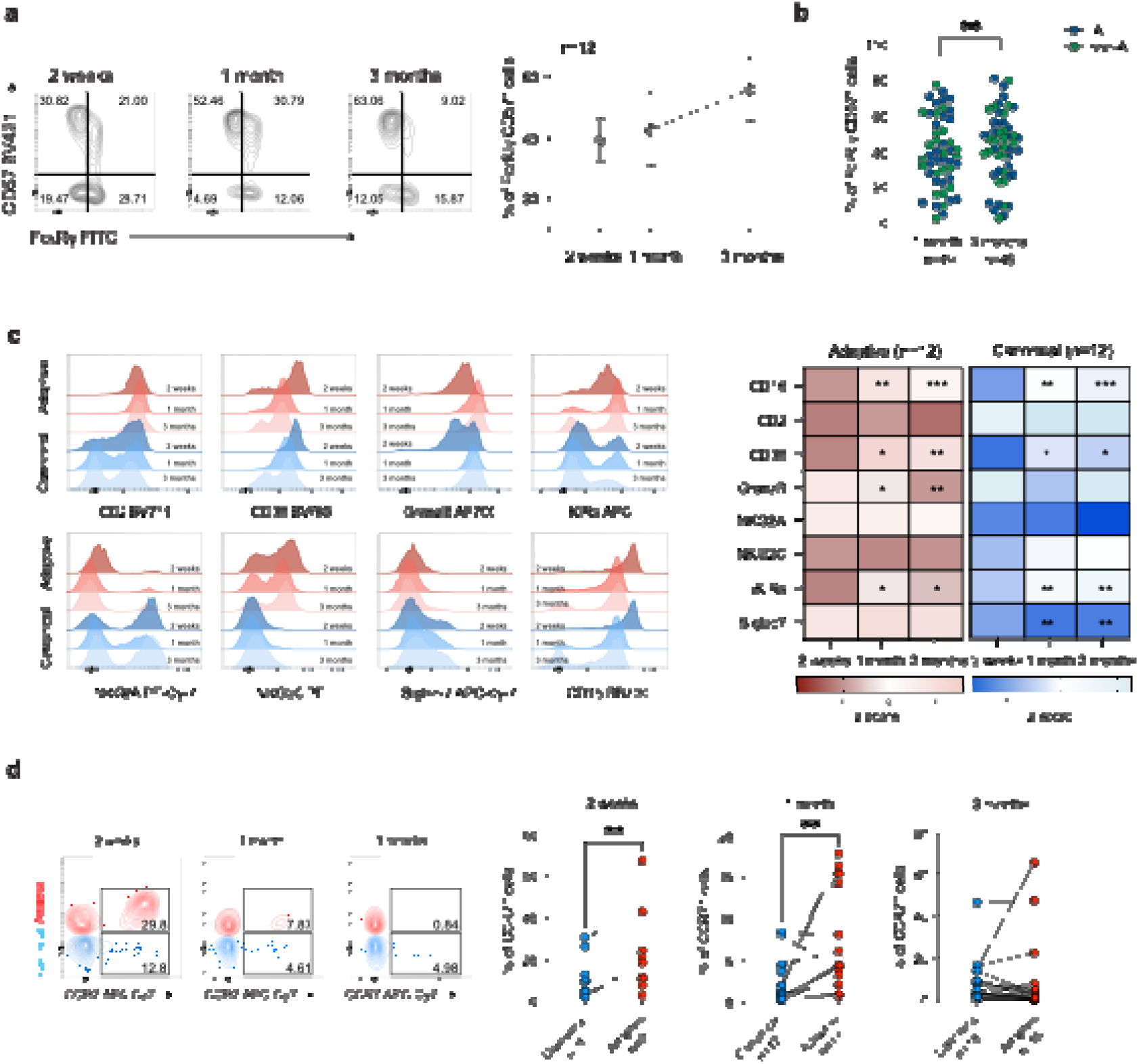
Progressive expansion of adaptive NK cells during acute HIV-1 infection. **(a)** Representative flow cytometry plots and longitudinal analysis of FcεRIγ^-^CD57^+^ adaptive NK cells in participants with samples available at all three timepoints (n = 12). **(b)** Summary data show the frequency of FcεRIγ^-^CD57^+^ adaptive NK cells at one and three months post-EDI in the whole cohort. Blue: subtype A; green: non-A subtypes. **(c)** Histograms and heatmap showing expression of indicated proteins in canonical and adaptive NK cells at two weeks, one month, and three months. Reported p values compare each timepoint with the 2-week timepoint. **(d)** Representative flow cytometry plots and summary data showing CCR7 expression on canonical (CD56^dim^NKG2C^-^) and adaptive (CD56^dim^NKG2C^+^) NK cells. Significance determined by two-tailed Wilcoxon matched-paired signed-rank test. *p<0.05, **p<0.01, ***p<0.001. Paired phenotyping data were available for 49 participants, as data at one month were missing for one participant. The longitudinal analysis included only a subset of participants with available samples at the 2-week timepoint. Due to limited sample availability, CCR7 analysis was performed in a subset of participants.

To determine whether adaptive NK cell expansion is reflected in ongoing proliferation, we assessed Ki-67 expression. The frequency of Ki-67^+^ cells within total NK cells was highest at two weeks and one month and progressively declined by three months (**Extended Data Fig. 3e,f**). However, adaptive NK cells had lower frequencies of Ki-67 cells than canonical NK cells at all three timepoints, suggesting that their enrichment during AHI may not reflect enhanced proliferation relative to canonical NK cells (**Extended Data Fig. 3g**).

We next asked whether adaptive NK cells retained functional features with the evolution of acute responses. Within FcεRIγ^-^ CD57^+^ adaptive and FcεRIγ^+^ CD57^+^ canonical NK cells, CD38, CD16 and iKIRs were highest at two weeks post-EDI, coinciding with peak viraemia, and declined by three months in both subsets (**Fig. 2c**). Granzyme B expression within adaptive NK cells increased over the same period, whereas CD2, which provides co-stimulation for CD16-dependent responses, showed no significant change (**Fig. 2c**). Siglec-7 increased in canonical NK cells but did not change within the adaptive subset, consistent with Siglec-7 loss being a stable feature of the adaptive phenotype ^21^ (**Fig. 2c**). Adaptive NK cells showed higher CCR7 expression than canonical NK cells at two weeks and one month, whereas no significant difference was detected at three months (**Fig. 2d**). Together, these data show that adaptive NK cells are already abundant during the earliest weeks of acute HIV-1 infection and are further enriched over the first three months, accompanied by increasing granzyme B and higher CCR7 expression confined to the first weeks of infection.

### Inflammatory cytokines and acute plasma augment adaptive NK cell expansion in vitro

Given the accumulation of adaptive NK cells during AHI, we sought to define signals and mechanisms that contribute to their early enrichment. We first considered CMV, a common co-infection in PLWH and a well-known driver of adaptive NK cell expansion ^5,21^. Plasma CMV DNA was undetectable in all participants tested at one month post-EDI (**Supplementary Table 3**), providing no evidence of systemic CMV reactivation at this timepoint.

We next measured 35 plasma biomarkers spanning cytokines, chemokines, angiogenesis, and vascular inflammation markers previously shown to be induced during hyperacute HIV-1 infection in the IAVI Protocol C cohort ^22^, to investigate whether these inflammatory signals contribute to the FcεRIγ adaptive NK cell expansion. Of the analytes measured, IL-12 and IL-15 correlated positively with adaptive NK cell frequency (z-scores, IL-12 r = 0.3782, p = 0.0104; IL-15 r = 0.4308, p = 0.0031) (**Fig. 3a**). We also calculated a composite score of the two cytokines, given that cytokines typically act in combination rather than in isolation. The composite IL-12/IL-15 z-score showed the strongest association (r = 0.4871, p = 0.0006; **Fig. 3a**). No significant correlations were found with other analytes and adaptive NK cell frequency (**Supplementary Table 4**).

**Figure 3.**
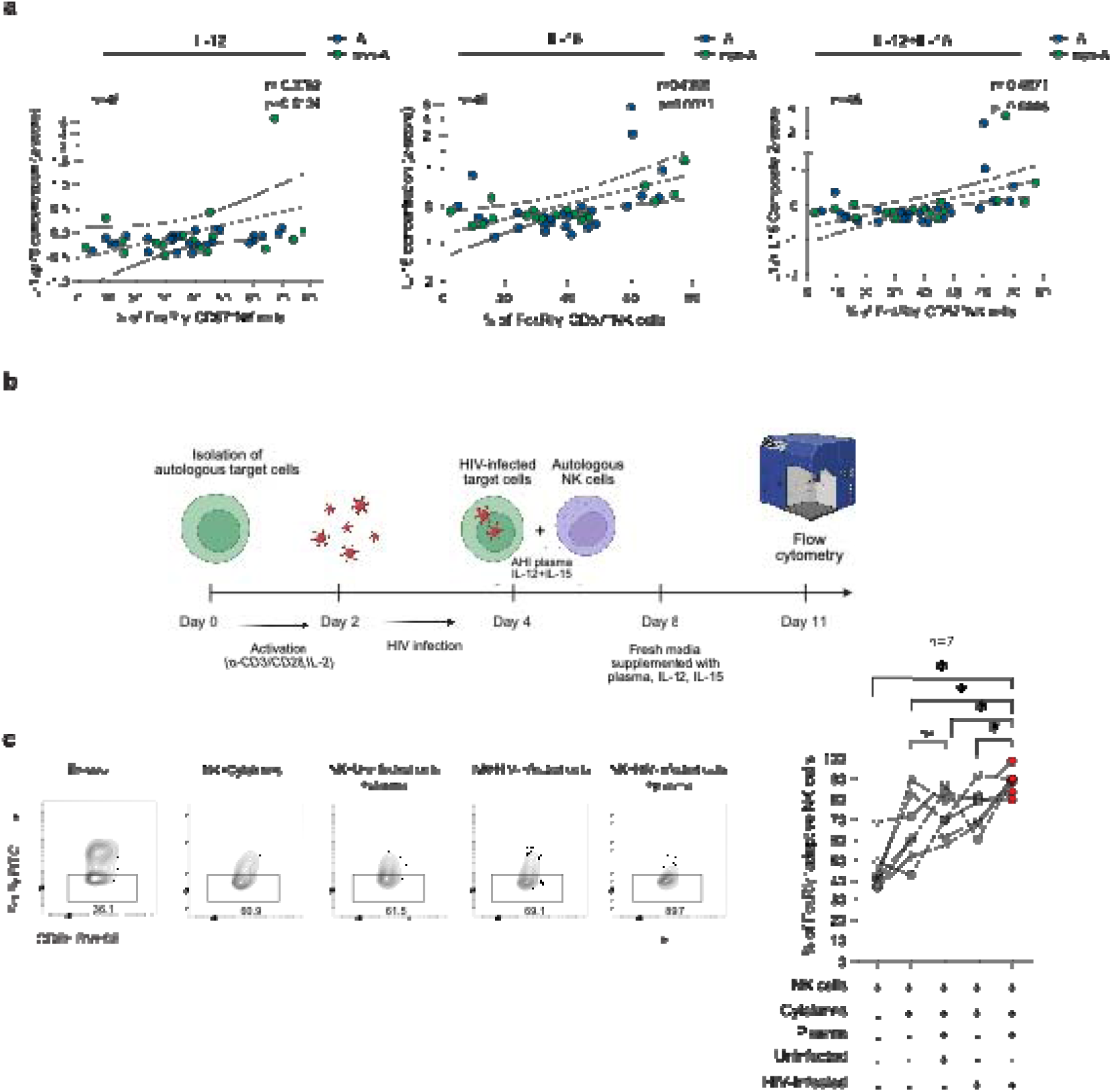
Inflammatory cytokines and acute plasma augment adaptive NK cell expansion. **(a)** Correlation between the percentage of FcεRIγ^-^CD57^+^CD56^dim^ NK cells and plasma concentration of IL-12, IL-15, and the combined IL12+IL15 composite z-score at one-month post-EDI. **(b)** Schematic graph showing the experimental process to set up activation, infection of target T cells, addition of acute plasma, and co-culture of autologous NK cells. **(c)** Representative example and summary data showing the proportion of FcεRIγ^-^ NK cells before and after co-culture with HIV-infected target CD4 T cells +/- plasma. Significance determined by Wilcoxon signed-rank test. *p<0.05. The non-parametric Spearman test was used for correlation analysis (two-tailed). The cytokine analysis was performed in 45 participants with available plasma samples. Expansion assays used n=7 independent ART-treated donors selected for low baseline frequencies of FcεRIγ CD57^+^ NK cells. All conditions were tested within each donor.

Adaptive NK cells expand in response to antibody-coated targets through CD16 engagement ^5,8^. We therefore investigated whether IL-12 and IL-15, together with autologous HIV-1-infected cells and acute-phase plasma, could augment FcεRIγ NK-cell expansion. Because acute Protocol C samples were limited and FcεRIγ NK cells were already expanded in these participants, we used samples from a different cohort of ART-treated PLWH with low baseline FcεRIγ NK cell frequencies to allow detection of expansion (**Supplementary Table 5)**. Purified NK cells were co-cultured with HIV-1-infected or uninfected target cells for 7-10 days in the presence of IL-12 and IL-15, with or without pooled plasma from Protocol C acute participants **(Fig. 3b)**. IL-12 and IL-15 increased the proportion of FcεRIγ adaptive NK cells relative to *ex vivo* levels (**Fig. 3c**). The addition of acute plasma and HIV-1-infected targets further increased this proportion, reaching significantly higher frequencies than cytokines alone, uninfected targets with plasma, or infected targets without plasma (**Fig. 3c**). Together, these findings suggest that IL-12 and IL-15, together with antibody-mediated recognition of HIV-1-infected cells, may contribute to adaptive NK cell enrichment during AHI.

### Adaptive NK cells mediate stronger antibody-dependent responses and greater degranulation against HIV-1-infected targets than canonical NK cells

Given the importance of ADCC in HIV-1 control ^23–26^, we evaluated ADCC capacity in our cohort by measuring the expression of CD107a and intracellular IFN-γ and TNF-α production after stimulation with Raji cells in the presence or absence of anti-CD20 antibody. Antibody-dependent IFN-γ NK cell responses were detectable from two weeks post-EDI and were maintained at one and three months, with no significant difference between the latter timepoints (**Fig. 4a,b**). These responses were preferentially mediated by FcεRIγ^-^ adaptive NK cells rather than canonical NK cells, with adaptive NK cells showing significantly stronger antibody-dependent IFN-γ responses than canonical NK cells at both one and three months (**Fig. 4c,d**). Neither total NK cell TNF-α production nor degranulation, measured by CD107a expression, differed significantly between one and three months post-EDI (**Extended Data Fig. 4a,b**). IFN-γ production also remained comparable between these timepoints within both adaptive and canonical NK cell populations (**Extended Data Fig. 4c**).

**Figure 4.**
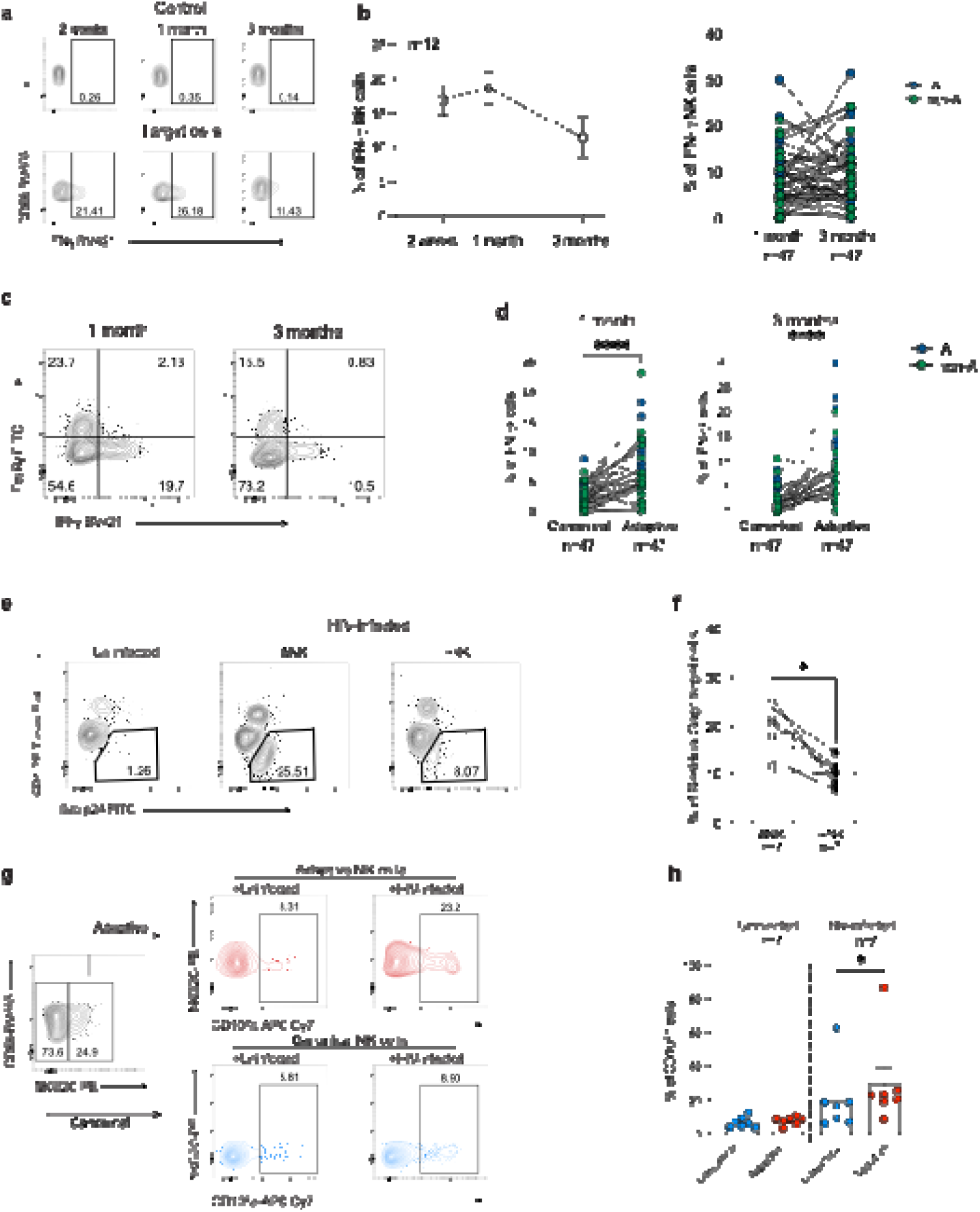
Longitudinal assessment of functional responses of adaptive NK cells during AHI. **(a)** Representative flow plots and **(b)** summary data for the frequency of IFN-γ-producing CD56^dim^ NK cells following 6-hour stimulation with antibody-coated Raji cells (Target cells) in all participants. Control = Uncoated cells. **(c)** Representative flow plots and **(d)** paired data showing IFN-γ production in FcεRI ^+^CD56^dim^ and FcεRIγ^-^CD56^dim^ NK cells at 1 month and 3 months post-EDI. **(e)** Example of the frequency of Gag p24^+^ cells in uninfected and HIV-1 infected conditions (with or without NK cell co-culture). **(f)** Summary data showing the frequency of Gag p24^+^ cells in the presence or absence of NK cells. **(g)** Representative flow cytometry plots of CD107a degranulation by NKG2C adaptive and NKG2C canonical NK cells after co-culture with autologous uninfected or HIV-1-infected CD4 T cells. **(h)** Paired analysis showing the percentage of CD107a-expressing cells within adaptive or canonical NK cells in response to uninfected or infected targets. Functional analyses included 47 participants with paired analysis after excluding data with high negative background responses. The killing assay comparing adaptive and canonical NK cell activity against autologous target cells was performed using samples from seven ART-treated participants. Significance determined by two-tailed Wilcoxon matched-paired signed-rank test. *p<0.05, **p<0.01, ****p < 0.0001.

Having shown that adaptive NK cells mediate stronger ADCC responses, we subsequently assessed their activity against autologous CD4 T cells infected with transmitted/founder (T/F) HIV-1 virus, a model that more closely reflects the viruses encountered during acute infection. Adaptive and canonical NK cells were defined using surface NKG2C expression, because intracellular FcεRIγ staining was not compatible with this assay; however, the majority of NKG2C CD56dim NK cells were FcεRIγ in these donors, supporting the use of NKG2C as a surface surrogate (**Extended Data Fig. 4d**). Compared with HIV-1-infected CD4 T cells cultured without NK cells, addition of autologous total NK cells reduced the frequency of p24 Gag CD4 T cells, consistent with NK cell-mediated elimination of infected targets **(Fig. 4e,f)**. While adaptive and canonical NK cells showed comparable degranulation in response to uninfected targets, adaptive NK cells exhibited greater CD107a expression in response to HIV-1-infected targets, suggesting preferential degranulation by the adaptive NK cell population **(Fig. 4g,h)**.

### Adaptive NK cell frequencies during the first month distinguish future controllers from non-controllers

We next asked whether adaptive NK cell expansion during AHI was associated with subsequent viral control. Using longitudinal plasma VL measurements, we identified seven participants who maintained persistently undetectable or low-level viraemia (<10,000 copies ml ¹) between 1.5 and 6 years post-EDI despite the absence of ART, allowing a single measurement above this threshold (n=7, future controllers, **Material and Methods**). We compared NK cell responses between the seven future controllers and twenty age-, sex- and subtype-matched non-controllers (VL>10,000 copies ml ¹) (**Fig. 5a, Supplementary Table 6**). This analysis was restricted to subtype A infection, as all identified controllers were living with subtype A and were compared with subtype A non-controllers to minimise potential confounding by infecting subtype.

**Figure 5.**
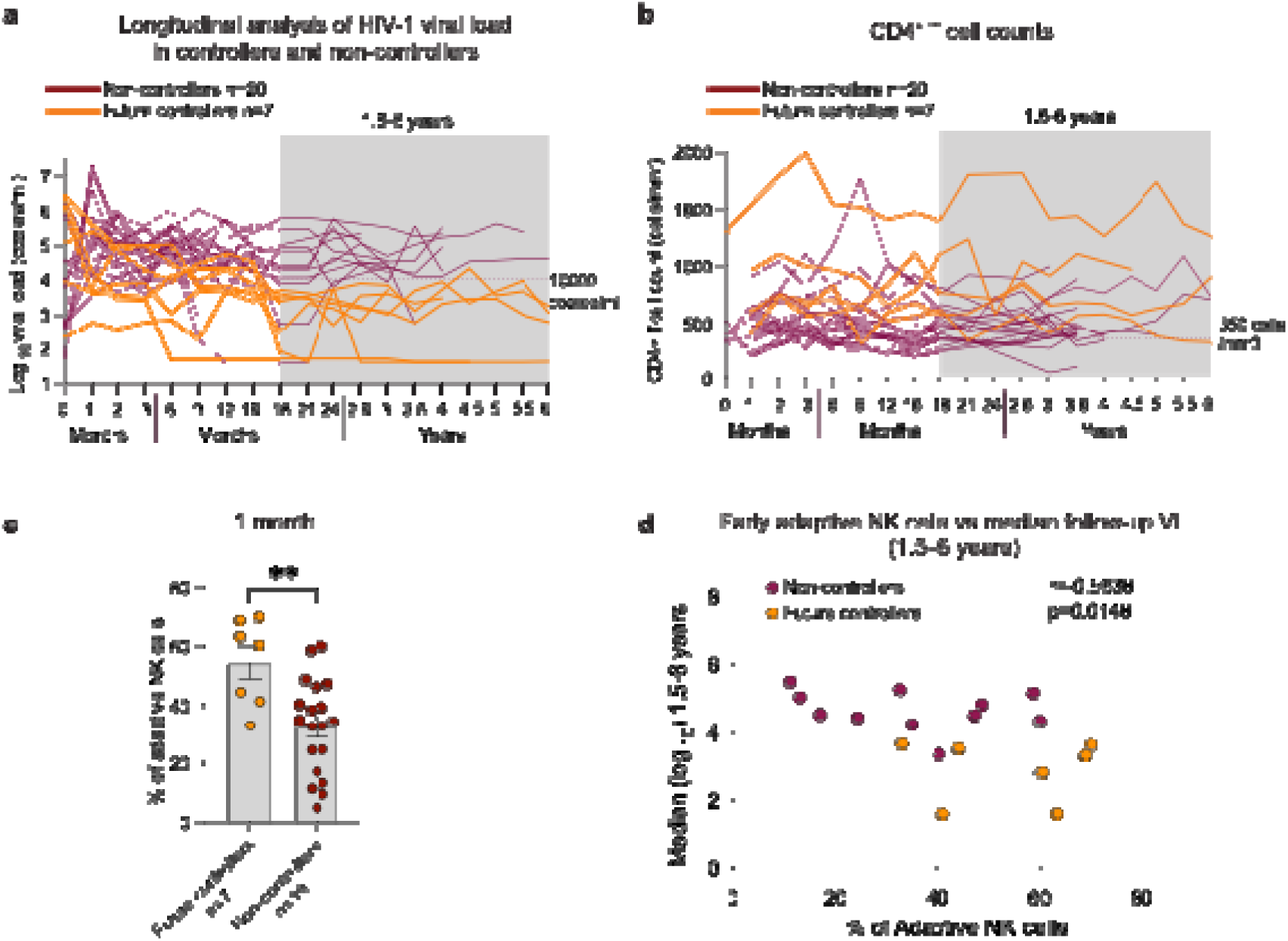
Adaptive NK cells during the first month of infection distinguish future controllers from non-controllers. **(a)** Log_10_ viral loads and **(b)** CD4^+^ T-cell counts of future controllers and non-controllers. Time-points used in defining virus control are highlighted in the shaded grey box. Future controllers maintain levels of HIV-1 viraemia <10,000 copies/ml within 1.5-6 years. **(c)** Frequency of adaptive NK cells at 1 month post-EDI in future controllers and non-controllers. **(d)** correlation analysis between the frequency of adaptive NK cells at 1 month and median log_10_ VL during untreated follow-up from 1.5-6 years post-EDI, shown for subtype A participants. Significance determined by two-tailed Mann-Whitney U test. The non-parametric Spearman test was used for correlation analysis (two-tailed).**p<0.01. One of the 20 subtype A non-controllers had data at three months but not at one month. Therefore, the analysis included 19 non-controllers at one month and 20 at three months.

Median log plasma viral load at one month was 4.22 (range 2.79 to 5.60) in future controllers and 5.06 (3.54 to 7.12) in non-controllers (**Supplementary Table 6**). Median CD4 T-cell counts were 599 and 518 cells/mm³ at one month, and 637 and 420 cells/mm³ at three months, respectively (**Supplementary Table 6**). During subsequent follow-up, counts in future controllers were generally above 350 cells/mm³, although some measurements fell below this threshold, with considerable variation between individuals (**Fig. 5b**). HLA-B alleles previously associated with slower disease progression ^16,27,28^ were present in 3 of 7 future controllers and 2 of 20 non-controllers **(Supplementary Table 6)**.

Future controllers had significantly higher adaptive NK cell frequencies than non-controllers at one month post-EDI (future controllers mean= 54.3%, non-controllers mean=32.4%, p = 0.005) **(Fig. 5c)**. At three months, adaptive NK cell frequencies remained higher in future controllers, although the difference did not reach significance (**Extended Data Fig. 5a**, p = 0.0733). Adaptive NK cell frequency at one month was also assessed against median log viral load across the same window, in participants with untreated measurements available. The two were inversely correlated in subtype A participants (r = −0.5638, p = 0.0148) **(Fig. 5d)**, but not across the full cohort (r = −0.2376, p = 0.1509) **(Extended Data Fig. 5b).**

Taken together, these data show that higher adaptive NK cell frequencies during the first month of HIV-1 infection distinguish participants who subsequently maintain viral control without treatment.

## Discussion

Durable control of HIV-1 without treatment has been attributed largely to adaptive immunity. Our findings suggest that differences in the NK cell compartment during acute HIV-1 infection may contribute to shaping subsequent viral control years later without ART. By studying participants before establishment of setpoint VL, we identified an association between early adaptive NK cell enrichment and subsequent untreated control among participants with subtype A infection. Their enhanced functional responses provide a possible explanation for this association, while the *in vitro* findings suggest that inflammatory signals and recognition of infected cells may help shape this early response.

Mature NK cell subsets with adaptive features (CD57^+^ FcεRIγ^-^NKG2C^+^) were abundant as early as two weeks post-EDI and became further enriched between one and three months, with granzyme B expression increasing over the first three months, consistent with cytotoxic potential. Ki-67 expression was consistently lower in adaptive than in canonical NK cells at all three timepoints. Adaptive NK cells have reduced mTOR activity and lower proliferative capacity than canonical NK cells, alongside elevated Bcl-2 and a metabolic profile associated with enhanced survival ^29,30^, and acquisition of the FcεRIγ^low^ phenotype is itself favoured by limited division ^30^. The increase in FcεRIγ NK cells may therefore reflect both preferential survival and loss of FcεRIγ expression by canonical NK cells. Adaptive NK cells expressed higher CCR7 levels than canonical NK cells at two weeks and one month, with no difference by three months. Transient CCR7 upregulation on NK cells occurs under inflammatory stimulation ^31^, which may confer access to secondary lymphoid tissue during peak viraemia, where NK cells have been shown to enter B cell follicles in SIV infection ^32^.

This early accumulation of adaptive NK cells was associated with plasma levels of IL-12 and IL-15, cytokines known to promote NK cell proliferation and survival through STAT4/5 activation ^33,34^, and that are sufficient to skew NK cells towards a memory phenotype *ex vivo* during HIV-1 infection ^35^. *In vitro*, the same cytokines drove adaptive NK cell expansion, with a further increase on exposure to HIV-1 infected targets in the presence of acute plasma but not with uninfected cells. This is consistent with engagement of antibody-coated infected targets through CD16 ^5,6^.

We did not detect CMV viraemia in our cohort at the timepoint tested, although localised reactivation in tissues cannot be excluded ^36^. Given near-universal seroprevalence in these populations ^37,38^, the observed expansion may reflect an increase in a pre-existing pool of CMV-associated adaptive NK cells ^21^ as described in chronic HIV-1 and HCMV co-infection^39^.

Functionally, adaptive NK cells exhibited enhanced antibody-dependent responses and stronger reactivity to HIV-1-infected target cells than canonical NK cells. The mechanisms through which adaptive NK cells recognise infected cells during acute infection remain to be fully defined but may involve altered classical HLA class I expression or recognition through the non-classical NKG2C–HLA-E axis. Evidence for the latter comes from studies showing that HLA-E-presented HIV-1 peptides activate NKG2C NK cells ^14^, and that, in people receiving long-term ART, sequence variation at NKG2C-contact residues within these epitopes is associated with reservoir size ^15^. Whether this pathway contributes to NK cell recognition during the first weeks of untreated infection, when antigen is abundant and classical HLA class I expression is downregulated, remains to be investigated.

Notably, participants who maintained undetectable or low viral loads (<10,000 copies/ml) between 1.5-6 years post-EDI without ART had a higher magnitude of adaptive NK cells at one month. These findings are consistent with the importance of these NK cell subsets in long-term viral control in elite and post-treatment controllers ^11,12,40^. Previous studies found that higher NKG2C CD57 NK cell frequencies were associated with faster viral suppression during ART ^41^ and identified early proliferative NK cells in two individuals who subsequently achieved spontaneous control ^42^. Our study extends these findings to control maintained in the absence of therapy, with the adaptive NK cell response quantified during the first month of infection and the outcome ascertained over up to six years of untreated follow-up. Their enhanced antibody-dependent activity and degranulation in response to infected cells with transmitted/founder virus provide a possible mechanism through which this early response could contribute to control of HIV-infected cells.

These observations have potential implications for vaccine design. The adaptive NK compartment can be assessed prior to immunisation, and may inform participant stratification or the interpretation of heterogeneity in vaccine efficacy, consistent with the association between adaptive NK cell frequencies and the magnitude of vaccine-induced responses reported in people living with HIV-1 ^43^. The compartment is also modifiable *in vitro*, raising the possibility that cytokine conditioning and Fc engagement could mobilise this population during priming.

There are limitations to this study. Our cohort comprised multiple HIV-1 subtypes; consistent with previous reports from this cohort ^16^, subtype A was associated with differences in viral load and CD4 T-cell trajectories. The controller analysis was restricted to subtype-A individuals, and the relationship between infecting subtype and adaptive NK cell responses warrants dedicated investigation. The observed heterogeneity in the dynamics of NK cell responses during AHI and the multi-regional nature of our cohort highlight the need to consider additional putative factors, including host genetics, as they relate to NK cell-mediated immunity ^44^. Moreover, due to the lack of pre-infection samples, it was not possible to establish whether the early enrichment of adaptive NK cells represented pre-existing populations. The sample size for the controller analysis is modest, reflecting the rarity of cohorts with sufficient untreated follow-up to define spontaneous control.

Despite these limitations, this cohort, sampled during acute infection and followed off ART for years, is unrepeatable under current treatment guidelines and will remain a reference point for the study of early immune events in untreated HIV-1 infection.

## Material and Methods

### Study design

Fifty ART-naïve participants from the IAVI Protocol C study were included in this study. IAVI Protocol C study was a prospective study conducted in Kenya, Rwanda, Uganda, and Zambia between 2006 and 2011 and has been previously described elsewhere ^16^. Samples were selected based on PBMC availability and the availability of matched specimens at both the 1- and 3-month timepoints. The samples were collected at three timepoints during AHI: two weeks (range: 9-19 days), one month (range: 21-42 days), and three months (range: 66-117 days) post-EDI. The estimated date of infection (EDI) was defined as 14 days before the date of the first positive p24 antigen test (with a negative antibody test), or 10 days before the date of the first PCR-positive test (with a negative antibody or p24 antigen detection) ^22,45^. Demographic and clinical data, including sex, ethnicity, HIV-1 diagnosis date, and transmission risk group, were obtained from the centralised curated IAVI Protocol C database.

All participants provided written informed consent. Research sites received approvals from respective country-specific ethics review boards: Kenya Medical Research Institute Ethical Review Committee; the Kenyatta National Hospital Ethical Review Committee of the University of Nairobi; the Rwanda National Ethics Committee, the Uganda Virus Research Institute Science and Ethics Committee; the Uganda National Council of Science and Technology; the University of Zambia Research Ethics Committee; and the Emory University Institutional Review Board ^16,46^. Further details on participants’ characteristics are included in **Supplementary Table 1.**

In addition to the Protocol C cohort, samples used for the *in vitro* adaptive NK cell expansion and co-culture assays (**Fig. 3b,c**, **Fig. 4e-h**) were obtained from an independent group of ART-treated PLWH, recruited at the Mortimer Market Centre for Sexual Health and HIV Research and the Royal Free Hospital (London, UK), with written informed consent from all donors. Donors were selected for low baseline FcεRIγ NK-cell frequencies, as acute Protocol C samples were limited. All donors were receiving suppressive ART with undetectable plasma HIV-1 RNA. Donor characteristics are summarised in **Supplementary Table 5**. The protocols for the human study were approved by the local Research Ethics Committee (REC)-Berkshire (REC 16/SC/0265). The study complied with all relevant ethical regulations for work with human participants and conformed to the Helsinki declaration principles and Good Clinical Practice (GCP) guidelines, and all subjects enrolled in the study provided written informed consent.

### HIV-1 subtype determination

The HIV-1 *env V1-V3* region (approximately 940 base pairs) was sequenced after extraction, reverse transcription, and polymerase chain reaction (PCR) amplification, as previously described ^22^. Generated sequences were aligned with HIV-1 subtype reference sequences from the Los Alamos HIV-1 sequence database (https://www.hiv.lanl.gov/content/index) and HIV-1 subtypes were phylogenetically determined as described ^47^.

### HIV-1 viral loads and CD4^+^ T-cell count measurements

Plasma viral load (RNA copies/ml) was measured in all participants using the Amplicor monitor assay (Roche Applied Science, Indianapolis, IN), as previously described ^48,49^. All viral load measurements were presented as log_10-_transformed values. CD4^+^ T-cell counts were collected at individual clinics using the FACScount System (Beckman Coulter Ltd., London, United Kingdom) ^50^. The number of participants with available viral loads or CD4^+^ T-cell counts (excluding counts after ART treatment) at each time-point presented in this study is listed in **Supplementary Table 2.**

### Definition of future controllers

Participants were classified as future controllers if untreated HIV-1 viral load remained below 10,000 copies/ml across available visits from 1.5-6 years post-EDI, allowing a single measurement above this threshold. Transient reductions below the threshold followed by viral rebound in most of the follow-up visits did not qualify as control. Only pre-ART measurements were included. One non-controller participant had low HIV-1 viral loads at three visits between months 9 and 21 but subsequently started ART; untreated follow-up within the 1.5-6-year window was therefore insufficient to confirm viral control. To minimise potential confounding by HIV-1 subtype, future controllers were compared with non-controllers with the same infecting subtype (subtype A).

### Sample preparation

PBMCs were thawed in a 37°C water bath. The thawed PBMCs were then transferred to a 50ml conical tube containing a pre-warmed complete RPMI medium (RPMI supplemented with penicillin-streptomycin, l-Glutamine, N-2-hydroxyethyl piperazine-N-2-ethane sulfonic acid (HEPES), non-essential amino acids, 2-Mercaptoethanol, and 10% FBS) and Benzonase (Dilution 1 in 2500 µl). Cells were then centrifuged for 10 minutes at 350 × g at room temperature and resuspended in a fresh complete RPMI medium. Cells were stained and counted using an Automated Cell Counter (BioRad, Hercules, California, USA). Cells were then resuspended at an appropriate cell concentration for each assay.

### Flow cytometric phenotypic assay

Cryopreserved PBMCs were thawed and rested for 1 h at 37 °C in a complete RPMI medium (as described above). Cells were then washed and plated in a 96-well plate at 0.5-1×10^6^ cells/well. Cells were then washed, resuspended in PBS, and surface stained at 4°C for 20 min with different combinations of antibodies in the presence of fixable live/dead stain (Invitrogen). Cells were then fixed and permeabilised for the detection of intracellular (Cytofix/Cytoperm™, BD Biosciences) or intranuclear (FoxP3 intranuclear staining buffer kit, eBioscience) antigens. Fixed cells were stained at room temperature for 30-45 min with different combinations of intracellular and intranuclear antibodies. Samples were acquired on a BD Fortessa X20 using BD FACSDiva8.0 (BD Biosciences), and subsequent data analysis was performed using FlowJo 10 (TreeStar). The full list of fluorochrome-conjugated antibodies and dilutions is included in **Supplementary Table 7**. The gating strategies used to define NK cell populations are provided in **Extended Data Fig. 2a**.

### Definition of adaptive NK cell populations

Adaptive NK cells were defined as FcεRIγ CD57 cells within the CD56^dim^ compartment for all *ex vivo* phenotypic analyses of the Protocol C cohort (**Fig. 2 and 3**). Unsupervised analysis of the same samples identified a corresponding population, metacluster 5, defined by NKG2C CD2 PLZF KIR Siglec-7 FcεRIγ CD57 (**Extended Data Fig. 3a-c**). This signature is more restrictive than the manual gating, so the two measures are related but not numerically equivalent. In the *in vitro* expansion assay, cells were identified by FcεRIγ alone (**Fig. 3c**). In the autologous target-cell assays, intracellular FcεRIγ staining was not compatible with the panel used, and adaptive and canonical NK cells were therefore defined by surface NKG2C expression (**Fig. 4g,h**); NKG2C CD56^dim^ NK cells in these donors were predominantly FcεRIγ (**Extended Data Fig. 4d**). The marker combination used is stated in each figure legend.

### NK cell functional assay

Purified PBMCs were thawed and rested for 1 h at 37 °C and 5% CO in a complete RPMI medium. Raji cells (target cells, the Burkitt lymphoma cell line Raji) were resuspended in complete RPMI at 5×10^5^/ml and incubated with 2.5 μg/ml of anti-human CD20 antibody (InvivoGen, France) or PBS as a negative control for 30 minutes at 37 °C. After 30 minutes, Raji cells were washed and resuspended to the desired concentration in complete RPMI. PBMCs from each participant were co-cultured with Raji cells at 5:1 ratio for 6 hrs at 37°C, in the presence of CD107a APC-H7 antibody (BD Biosciences, dilution 1 in 200 μl). After 1 h, GolgiStop (containing Monensin, 2 μmol/ml) and GolgiPlug (containing brefeldin A, 10 μg/ml) (BD Biosciences) were added to inhibit protein transport. After stimulation, cells were washed and surface stained at 4°C for 20 min in the presence of fixable live/dead stain (Invitrogen). Cells were then fixed and permeabilised (CytoFix/CytoPerm; BD Biosciences), followed by intracellular cytokine staining with IFN-γ BV421 (BioLegend, dilution 1 in 100), and TNF-α BV711 (BioLegend, dilution 1 in 50). Samples were acquired on a BD Fortessa X20 using BD FACSDiva8.0 (BD Biosciences), and data were analysed using FlowJo 10 (TreeStar).

### Preparation of HIV-1 infected target cells

CD4^+^ T cells were isolated from frozen PBMCs using CD4^+^ T-cell Isolation Kit (Miltenyi Biotec, Catalog # 130-096-533) as per the manufacturer’s instructions. CD4^+^ T cells were activated using T-cell Activation Kit (Thermofisher, Catalog # 11161D) in complete RPMI medium containing 10 ng/mL IL-2 (R&D Systems, Minneapolis, MN, Cat#202-IL-500) for 48 hours. The activated CD4^+^ T cells were then resuspended to 5 million/mL in complete RPMI medium + 10ng/mL IL-2, and then 200 µl of cells was added to each well of the 96 well plate (1 million/well). After two days of activation, the cells were removed from the plate, washed in media, and infected with T/F virus (CH58) at 1 million cells/well in media + 10ng/mL IL-2. The cells were spinoculated at 800 g for two hours at 20-25°C. Upon completion of the spinfection, cells were resuspended at a cell density of 2×10^6^/mL in RPMI-complete supplemented with 10 ng/mL IL-2. Uninfected, activated CD4^+^ T cells were incubated at 37°C incubator during the process of infection. Cells were then plated in 96-well plates and incubated at 37°C for 2-3 days.

### NK cell elimination assay

Autologous NK cells were enriched from frozen PBMCs using NK Cell Isolation Kit (Miltenyi Biotec, Catalog # 130-092-657) as per the manufacturer’s instructions (>80% purity and viability). NK cells were rested overnight in complete RPMI medium containing 50 ng/mL IL-2 (R&D Systems, Minneapolis, MN, Catalog#202-IL-500). Uninfected and infected CD4^+^ T cells were prepared as described above. In 96 well round bottom plates, NK cells were co-cultured with uninfected and infected target T cells at 1:1 effector-to-target ratio (E:T) for four hours at 37°C. After incubation, cells were surface stained at 4°C for 20 min with different combinations of antibodies (**Supplementary Table 7**) in the presence of fixable live/dead stain (Invitrogen). Cells were then fixed and permeabilised for the detection of intracellular (Cytofix/Cytoperm™, BD Biosciences). Fixed cells were stained for the HIV-1 Gag protein using anti-Gag p24-RD1 (Beckman Coulter, Catalog #6604667, dilution 1 in 50) at room temperature for 30 min.

### NK cell expansion assay

For the expansion assay, HIV-1 infected target T cells (or mock-infected cells) were prepared as described above. Autologous NK cells were enriched from frozen PBMCs using the NK Cell Isolation Kit (Miltenyi Biotec, Catalog # 130-092-657) as per the manufacturer’s instructions. Target T cells and NK cells were co-cultured for 7-10 days in the presence of recombinant human IL-12 and IL-15 (100ng/ml). The culture contained pooled plasma from 18 acutely infected participants from the IAVI Protocol C cohort, collected mainly within the first three months post-EDI. During the culture, half of the culture media was removed and replenished with fresh media. At the end of co-culture, the proportions of adaptive NK cells were measured using flow cytometry.

### Quantification of CMV DNA

CMV DNA was quantified using an in-house qPCR assay based on previously published work ^51^. Plasma DNA containing cell-free CMV was extracted using Qiazol (Qiagen) and chloroform. Briefly, 100 ul of plasma was incubated with 1 ml of Qiazol at room temperature for 10 minutes. Then, 200 ul of chloroform was added and shaken for 15 seconds. The sample was centrifuged at 12000 g for 15 minutes at 4°C, after which the aqueous phase was removed. Pure ethanol (0.3 ml) was added to the DNA-containing interphase and phenol phase and mixed by inversion. The samples were incubated at room temperature for 2-3 minutes and centrifuged at 2000 g for 2 minutes at 4°C to sediment DNA. The DNA pellet was washed three times with 1 ml sodium citrate, each time with a 30-minute incubation at room temperature (during which mixing by inversion was done every 5 minutes) followed by centrifugation at 2000xg for 5 minutes at 4°C. Then, 2 ml of 75% ethanol was added to the DNA pellet and incubated at room temperature for 20 minutes. The sample was centrifuged at 2000 g for 5 minutes at 4°C, and the ethanol supernatant was removed. The DNA pellet was air-dried for 5-15 minutes before re-dissolving in 50 ul of 8mM NaOH. The reaction mix consisted of 10 ul Maxima qPCR Master Mix (2X), 0.6 ul of each primer (final concentration of 0.3 uM), 0.4 ul of the probe (final concentration of 0.2 uM), 5.9 ul of nuclease-free water, and 2.5 ul of DNA template in a reaction volume of 20 ul. The cycling program was 1 cycle of 95°C for ten minutes (enzyme activation) followed by 50 cycles of 95°C for 15 seconds and 60°C for 60 seconds. PCR efficiency ranged from 89-97% across seven runs, with an inter-assay CV of 16% based on the internal controls.

### Measurement of cytokines and chemokines

Cytokine concentrations in plasma were measured using the MesoScaleDiscovery (MSD) multisport assay (MesoScaleDiagnostics, LLC) according to the manufacturer’s instructions ^22^. The values for analyte concentration, including respective lower (LLOQ) and upper (ULOQ) limits of quantification, were calculated from standard curves. Composite cytokine z-scores were generated using the following method; Cytokine concentrations were log10-transformed and then standardised across all participants as z-scores using the cohort mean and standard deviation (SD) of the log-transformed values. The IL12+IL15 composite z-score was calculated as the mean of the IL-12 and IL-15 z-scores. A full list of biomarkers measured in this study is listed in **Supplementary Table 4.**

### High-dimensional data analysis

To evaluate the co-expression of markers at a single-cell level and to identify various cell clusters with similar phenotypic profiles, unsupervised multidimensional analysis was performed using different algorithms in Cytobank (https://www.cytobank.org). Concatenated files were used to evaluate the overall immune cell landscape in different groups. Cells were manually gated for lymphocytes, singlets, live cells, and CD3^-^CD7^+^CD56^dim^CD16^+^/^-^, and then subjected to viSNE analysis. The viSNE clustering analysis was performed on 10 parameters (FcεRIγ, NKG2C, PLZF, NKG2A, iKIRs, Siglec-7, GranB, CD57, CD2, and CD38). Equal events of 20,000 cells were selected across all samples. After vi-SNE analysis, FlowSOM was performed using algorithm Self-Organising Maps (SOMs) to define different clusters based on selected markers (FcεRIγ, NKG2C, PLZF, NKG2A, iKIRs, Siglec-7, GranB, CD57, CD2, and CD38) and reveal related phenotypic clusters ^52^. The number of metaclusters was set to 7-10, the number of clusters was set to 100, and the size of clusters was set to relative with 15 pixels as Max relative size (Cytobank default).

### Statistical analyses

Prism 8 (GraphPad Software) was used for statistical analysis as follows: The Mann-Whitney U test was used for single comparisons of independent groups; the Wilcoxon matched-pairs signed-rank test was used to compare two paired groups. The Spearman test was used for correlation analysis. viSNE and FlowSOM analyses were performed using Cytobank (https://www.cytobank.org). Statistical significance was indicated in the figures as *p<0.05, **p<0.01, ***p<0.001, and ****p<0.0001, and all tests were two-tailed.

## Data availability

All the data presented in this study are available in the published article and summarised in the corresponding tables, figures, and supplemental materials. Source data will be provided with the paper.

## Acknowledgements

The authors are grateful to all the clinic staff and study participants. We also thank IAVI for supporting HIV-1 research studies and capacity building initiatives in Kenya, Uganda, Rwanda, and Zambia. IAVI’s work is made possible by generous support from many donors, including the Bill & Melinda Gates Foundation; the Ministry of Foreign Affairs of Denmark; Irish Aid; the Ministry of Finance of Japan; the Ministry of Foreign Affairs of The Netherlands; the Norwegian Agency for Development Cooperation (NORAD); and the United States Agency for International Development (USAID). The full list of IAVI donors is available at www.iavi.org. The contents of this manuscript are the responsibility of the authors and do not necessarily reflect the views of USAID or the US Government.

## Funding

Saudi Ministry of Education Scholarship FG-350441 and KAUST The Ibn Rushd Postdoctoral Fellowship (AA); British HIV Association research award (DP); AIDS International Research Project of Centre for AIDS Research (SRJ); Kumamoto University (SRJ with Professor Masafumi Takiguchi); National Institutes of Health grant award R01AI155182 (DP); Swedish Research Council, 2016-01417 and 2020-06262 (JE) and Swedish Society for Medical Research SA-2016 (JE).

## Author contributions

A.A. performed experiments, acquisition of data, analysis, and drafting of the manuscript; A.S.H., J.H. and A.H. performed experiments and contributed to the acquisition of data and analysis. J.G., M.A.P, W.K., E.K., and E.J.S. contributed to patient recruitment, clinical sample processing and curating clinical data. A.S.H, J.G., and M.A.P. contributed to data interpretation and critical editing of the manuscript. J.E., E.J.S., D.P. and S.R.J. contributed to the conception and design of the study, data interpretation, critical revision of the manuscript and study supervision.

## Ethics declarations Competing interests

The authors declare no competing interests.

## Notes

### Competing Interest Statement

The authors have declared no competing interest.

### Summary of Updates

This version reports a revised analysis with a different focus from version 1, centred on NK cell responses during acute infection and their association with subsequent viral control. Some analyses from version 1 do not form part of this version, and additional experiments have been added.

