## SUPPLEMENTARY MATERIALS for "Early enrichment of adaptive NK cells during acute HIV-1 infection is associated with subsequent durable viral control"

Extended data figures and supplementary tables

**Extended Data Figure 1**. HIV-1 viral load and CD4⁺ T-cell counts over time in participants with subtype A vs non-A infection.

**Extended Data Figure 2.** Distribution of NK cell subsets during AHI.

**Extended Data Figure 3.** The landscape of CD56^dim^ NK cells over the course of AHI.

**Extended Data Figure 4.** TNF-α production and CD107a expression by total NK cells

**Extended Data Figure 5.** Comparison of adaptive NK cells between future controllers vs non-controllers at 3 months post-EDI

**Supplementary Table 1.** Cohort Demographics and Clinical Characteristics.

**Supplementary Table 2.** Comparisons of HIV VL and CD4+ T-cell counts between subtype A and non-A subtypes.

**Supplementary Table 3.** Detection of CMV viraemia during AHI.

**Supplementary Table 4.** Correlations between adaptive NK cell frequencies at 1 month and inflammatory biomarkers measured in this study.

**Supplementary Table 5.** Characteristics of ART-treated participants. Samples from these participants were used for in vitro co-culture assays.

**Supplementary Table 6.** Clinical characteristics of future controllers and non-controllers.

**Supplementary Table 7.** Antibodies used in flow cytometric analysis.

**Extended data figures**

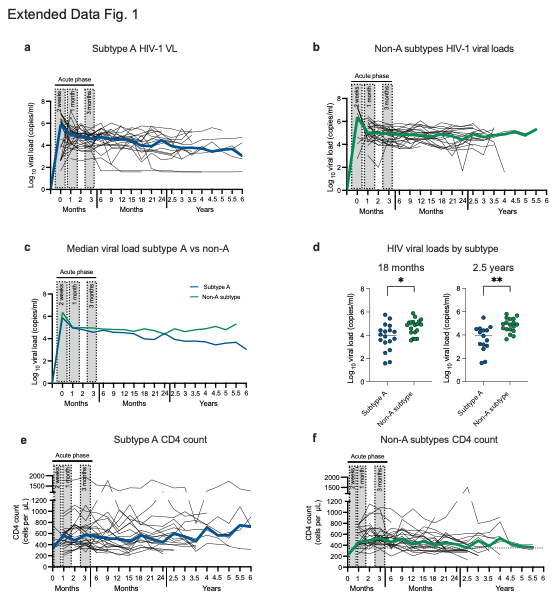

**Extended Data Figure 1. HIV-1 viral load and CD4⁺ T-cell counts over time in participants with subtype A vs non-A infection.**

**(a)** Log_10_ VL for subtype A and **(b)** non-A subtype participants following the onset of detectable plasma viraemia and up to six years. All measurements after the initiation of ART were excluded from the analysis. The thickened line represents the median log_10_ viral load. The timepoints sampled in this study are highlighted in the grey-shaded box (1^st^ time point = two weeks; 2^nd^ time point = one month; 3^rd^ time point = three months). **(c)** Comparison of median log_10_ viral load for participants with subtype A and non-A subtype infection. **(d)** HIV viral loads by subtype at 18 months and 2.5 years. Viral load measurements after ART initiation are excluded. **(e)** CD4^+^ T-cell counts for subtype A and **(f)** non-A subtype participants following the onset of detectable plasma viraemia and up to 3.5 years. The timepoints sampled in this study are highlighted in the grey-shaded box.

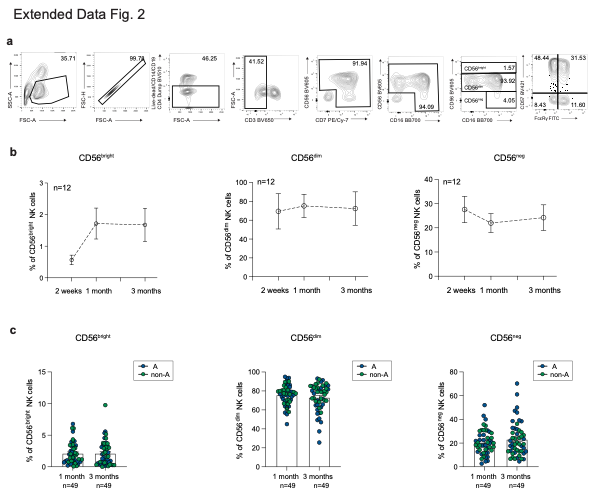

**Extended Data Figure 2. Distribution of NK cell subsets during AHI.**

**(a)** Example of gating strategy used to identify NK cell subsets. Cells were gated on a single lymphocyte population by side and forward side scatter, followed by live cells gating by excluding dead, CD4^+^, CD19^+^, and CD14^+^ cells. Following gating on live CD3^-^ T cells, cells were gated on CD56^+^/^-^ and CD7^+^/^-^. Total NK cells were then identified by gating in CD16^+^/^-^ and CD56^+^/^-^. **(b)** Percentage of NK cell subsets (CD56^bright^, CD56^dim^, and CD56^neg^ NK cells) at two weeks, 1 month, and three months post-EDI in a subset of participants (n=12). **(c)** Percentage of CD56^bright^, CD56^dim^, and CD56^neg^ NK cells in all participants at one and three months. Significance determined by two-tailed Mann-Whitney U test or Wilcoxon matched-pairs signed rank test. *p<0.05, **p<0.01.

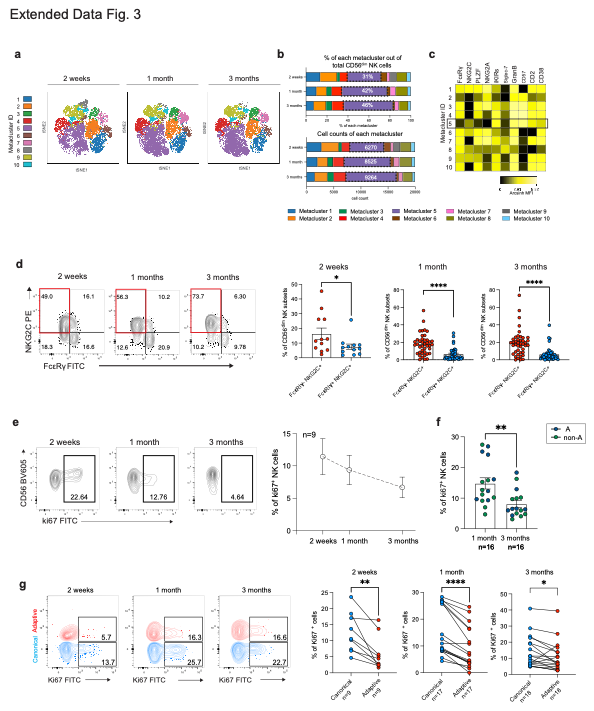

**Extended Data Figure 3. The landscape of CD56^dim^ NK cells over the course of AHI.**

**(a)** vi-SNE projections of concatenated flow cytometry data for CD56^dim^ NK cells from samples collected at two weeks, one month or three months post-infection. Each point on the high-dimensional mapping represents an individual cell, and meta-clusters are colour-coded. **(b)** Bar chart representing the cell counts and proportion of each FlowSOM meta-cluster out of the total CD56^dim^ NK cells. **(c)** Expression intensity heatmap of the indicated markers (FcεRIγ, NKG2C, PLZF, NKG2A, iKIRs, Siglec-7, GranB, CD57, CD2, and CD38) for each FlowSOM meta-cluster. **(d)** Representative flow cytometry plots and summary data showing frequencies of FcεRIγ⁺NKG2C⁺CD56^dim^ and FcεRIγ^-^NKG2C⁺CD56^dim^ NK cells. **(e,f)** Representative flow plots and summary analysis of the percentage of Ki-67^+^NK cells within total NK cells. **(g)** Representative plots and summary analysis of the percentage of Ki-67^+^NK cells within NKG2C^-^CD56^dim^ canonical NK cells and NKG2C^+^CD56^dim^adaptive NK cells. two-tailed Mann-Whitney U test or Wilcoxon matched-pairs signed rank test. *p < 0.05, **p < 0.01, ****p < 0.0001.

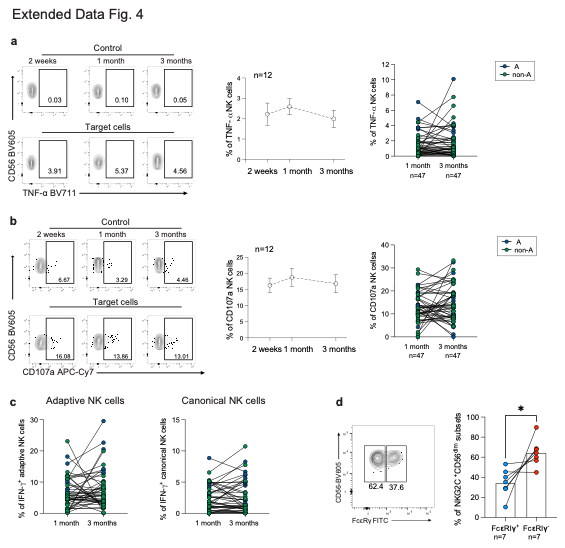

**Extended Data Figure 4. TNF-α production and CD107a expression by total NK cells**

**(a**) Representative flow plots and summary data for the frequency of TNFα- or **(b)** CD107a-producing CD56^dim^ NK cells following 6 hours of stimulation with antibody-coated Raji cells (Target cells). Control = the antibody-uncoated condition (the negative control of the assay). **(c)** Longitudinal comparison of antibody-dependent IFN-γ responses in adaptive (FcεRIγ⁻CD56^dim^) and canonical (FcεRIγ⁺CD56^dim^) NK cells at one and three months post-EDI. **(d)** Representative flow cytometry plots and paired data showing the percentage of FcεRIγ⁻ and FcεRIγ⁺ cells within NKG2C⁺ CD56^dim^ NK cells. Significance determined by two-sided paired Wilcoxon signed-rank tests. *p<0.05.

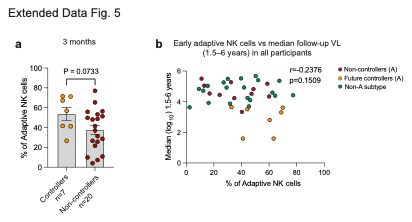

**Extended Data Figure 5. Comparison of adaptive NK cells between future controllers vs non-controllers at 3 months post-EDI.**

**(a)** Frequency of FcεRIγ⁻CD57⁺ adaptive NK cells at 3 months post-EDI in future controllers and non-controllers. **(b)** Association between FcεRIγ⁻CD57⁺ adaptive NK cell frequencies at one month and median log₁₀ VL during untreated follow-up from 1.5-6 years post-EDI, shown for the full cohort.

**Supplementary Table 1. Cohort Demographics and Clinical Characteristics.**

|  |  |  | **Subtype A** | **Non-A subtypes** |
| --- | --- | --- | --- | --- |
| **Group size** | No of participants (n) | Two weeks | 9 | 3 |
|  |  | One month | 27 | 23 |
|  |  | Three months | 27 | 23 |
| **HIV-1 subtypes** | HIV-1  subtypes (n, %) | A | 27 (100.0) | 0 (0.0) |
|  |  | C | 0 (0.0) | 14 (60.9) |
|  |  | D | 0 (0.0) | 9 (39.1) |
| **Enrolment** | Days post-EDI ^a^ | Two weeks, median (range) | 17 (9-19) | 18 (15-18) |
|  |  | One month, median (range) | 30 (21-42) | 36 (25-42) |
|  |  | Three months,  median (range) | 87 (66-110) | 89 (75-117) |
| **Recruitment**  **clinical site** | Research centre, country | Kigali, Rwanda, n (%) | 10 (37.0) | 0 (0.0) |
|  |  | Kilifi, Kenya, n (%) | 17 (63.0) | 4 (17.4) |
|  |  | Masaka, Uganda, n (%) | 0 (0.0) | 7 (30.4) |
|  |  | Lusaka, Zambia, n (%) | 0 (0.0) | 12 (52.2) |
| **Risk groups** | Age | Age (years), median (range) | 26 (19-52) | 33 (17-51) |
|  | Sex | female:male (n) | 3:24 | 5:18 |
|  | Risk group | Discordant couple (DC),  n (%) | 10 (37) | 19 (82.6) |
|  |  | men who have sex with men (MSM), n (%) | 15 (55.6) | 4 (17.4) |
|  |  | Other heterosexual, n (%) | 2 (7.4) | 0 (0.0) |
| **HIV-1 parameters** | Two weeks | Log_10_ HIV-1 viral load, median (range) | 6.38 (3.83-7.30) | 5.54 (5.14-7.43) |
|  |  | CD4^+^ T-cell count,  median (range) | 346 (189-1295) | 621 (478-720) |
|  | One month | Log_10_ HIV-1 viral load, median (range) | 5.01 (2.78-7.12) | 5.02 (3.04-6.10) |
|  |  | CD4^+^ T-cell count,  median (range) | 553 (189-1759) | 524 (245-828) |
|  | Three months | Log_10_ HIV-1 viral load, median (range) | 4.87 (2.79-6.03) | 4.82 (3.9-5.7) |
|  |  | CD4^+^ T-cell count,  median (range) | 466 (267-2000) | 503 (322- 1090) |
|  | Set-point viral load ^b^ | Log_10_ set-point,  median (range) | 4.52 (1.66-5.69) | 4.65 (3.3-5.77) |
| **Acute retroviral syndrome (ARS) symptoms** |  | Night Sweats, n (%) | 12 (44.4) | 4 (17.4) |
|  |  | Myalgia, n (%) | 18 (66.6) | 6 (26.1) |
|  |  | Fatigue, n (%) | 18 (66.6) | 2 (8.7) |
|  |  | Skin Rash, n (%) | 2 (7.4) | 0 (0.0) |
|  |  | Oral ulcers, n (%) | 6 (22.2) | 2 (8.7) |
|  |  | Pharyngitis, n (%) | 10 (37.0) | 0 (0.0) |
|  |  | Lymphadenopathy, n (%) | 10 (37.0) | 2 (8.7) |
|  |  | Diarrhoea, n (%) | 6 (22.2) | 5 (21.7) |
|  |  | Anorexia, n (%) | 15 (55.6) | 19 (82.6) |

^a^ The estimated date of infection (EDI) was defined as the date of a high-risk exposure incident, midpoint between the last negative and the first

HIV-1 antibody positive test, 10 days prior to the first positive HIV-1 PCR test, or 14 days prior to the first positive p24 antigen test.

^b^ Set-point viral load (VL) was defined according to the calculations for the Protocol C dataset. Data was missing for *n*=8 subtype A, *n*=7 non-A subtype.

**Supplementary Table 2. Comparisons of HIV VL and CD4^+^ T-cell counts between subtype A and non-A subtypes.**

|  |  | HIV Viral load | | | | | CD4 T-cell counts | | | | |
| --- | --- | --- | --- | --- | --- | --- | --- | --- | --- | --- | --- |
|  |  | Number of participants (n) | | Median HIV VL | |  | Number of participants (n) | | Median CD4 counts | |  |
| Timepoint | Days post-EDI | Subtype A | Non-A subtypes | Subtype A | Non-A subtypes | P value_ | Subtype A | Non-A subtypes | Subtype A | Non-A subtypes | P value |
| Month 0 | 0-20 | 23 | 8 | 5.88 | 6.36 | ns | 7 | 1 | 346 | 214 | ns |
| Month 1 | 21-45 | 25 | 23 | 4.95 | 4.99 | ns | 24 | 21 | 560 | 453 | ns |
| Month 2 | 46-76 | 24 | 23 | 4.81 | 4.98 | ns | 23 | 23 | 479 | 490 | ns |
| Month 3 | 77-136 | 24 | 22 | 4.58 | 4.96 | ns | 24 | 21 | 574 | 528 | ns |
| Month 6 | 137-228 | 24 | 23 | 4.75 | 4.86 | ns | 26 | 23 | 542 | 460 | ns |
| Month 9 | 229-319 | 22 | 21 | 4.57 | 4.82 | ns | 23 | 21 | 510 | 517 | ns |
| Month 12 _ | 320-410 | 20 | 20 | 4.55 | 4.81 | ns | 20 | 20 | 505 | 422 | ns |
| Month 15 | 411-502 | 21 | 20 | 4.44 | 4.66 | ns | 22 | 22 | 452 | 470 | ns |
| Month 18 | 503-593 | 18 | 19 | 3.98 | 4.88 | **0.015** | 18 | 22 | 567 | 408 | **0.051** |
| Month 21 | 594-684 | 17 | 15 | 3.93 | 4.72 | ns | 16 | 18 | 501 | 441 | ns |
| Month 24 | 685-821 | 9 | 8 | 4.46 | 4.42 | ns | 8 | 7 | 445 | 418 | ns |
| Year 2.5 | 822-1,004 | 15 | 18 | 3.93 | 4.89 | **0.001** | 14 | 18 | 603 | 353 | **0.003** |
| Year 3 | 1,005-1,187 | 14 | 12 | 3.77 | 4.63 | **0.060** | 14 | 11 | 543 | 483 | **0.095** |
| Year 3.5 | 1,188-1,369 | 12 | 7 | 3.78 | 4.79 | ns | 10 | 8 | 462 | 389 | ns |
| Year 4 | 1,370-1,552 | 7 | 4 | 3.71 | 4.88 | ns | 6 | 6 | 710 | 527 | ns |
| Year 4.5 | 1,553-1,735 | 5 | 2 | 3.45 | 5.10 | ns | 5 | 2 | 566 | 423 | ns |
| Year 5 | 1,736-1,917 | 6 | 2 | 3.60 | 4.81 | ns | 6 | 2 | 568 | 379 | ns |
| Year 5.5 | 1,918-2,100 | 5 | 1 | 3.67 | 5.30 | ns | 5 | 2 | 748 | 361 | ns |
| Year 6 | 2,101-2,282 | 5 | 0 | 3.04 |  |  | 5 | 0 | 725 |  |  |

* Two-sided, unpaired Mann–Whitney U. The tests use individual values, not the reported medians.

**Supplementary Table 3. Detection of CMV viraemia during AHI.**

| **Subtype** | **PID** | **Plasma CMV DNA (copies/ml)** |
| --- | --- | --- |
| Subtype A | PID1 | Negative |
| Subtype A | PID2 | Negative |
| Subtype A | PID3 | Negative |
| Subtype A | PID4 | Negative |
| Subtype A | PID5 | Negative |
| Subtype A | PID6 | Negative |
| Subtype A | PID7 | Negative |
| Subtype A | PID8 | Negative |
| Subtype A | PID9 | Negative |
| Subtype A | PID10 | Negative |
| Subtype A | PID11 | Negative |
| Subtype A | PID12 | Negative |
| Subtype A | PID13 | Negative |
| Subtype A | PID14 | Negative |
| Subtype A | PID15 | Negative |
| Subtype A | PID16 | Negative |
| Subtype A | PID17 | Negative |
| Subtype A | PID18 | Negative |
| Subtype A | PID19 | Negative |
| Subtype A | PID20 | Negative |
| Subtype A | PID21 | Negative |
| Subtype A | PID22 | Negative |
| Subtype A | PID23 | Negative |
| Subtype A | PID24 | Negative |
| Subtype A | PID25 | Negative |
| Subtype A | PID26 | Negative |
| Subtype A | PID27 | Negative |
| Subtype C | PID28 | Negative |
| Subtype C | PID29 | Negative |
| Subtype C | PID30 | Negative |
| Subtype C | PID31 | Negative |
| Subtype C | PID32 | Negative |
| Subtype C | PID33 | Negative |
| Subtype C | PID34 | Negative |
| Subtype C | PID35 | Negative |
| Subtype C | PID36 | Negative |
| Subtype C | PID37 | Negative |
| Subtype C | PID38 | Negative |
| Subtype C | PID39 | Negative |
| Subtype C | PID40 | Negative |
| Subtype C | PID41 | Negative |
| Subtype D | PID42 | Negative |
| Subtype D | PID43 | Negative |
| Subtype D | PID44 | Negative |
| Subtype D | PID45 | Negative |
| Subtype D | PID46 | Negative |
| Subtype D | PID47 | Negative |
| Subtype D | PID48 | Negative |
| Subtype D | PID49 | Negative |
| Subtype D | PID50 | Negative |

**Supplementary Table 4. Correlations between adaptive NK cell frequencies at 1 month and inflammatory biomarkers measured in this study.**

| Panel | Abbreviation | Full name | Spearman r | P value |
| --- | --- | --- | --- | --- |
| Cytokine | GM-CSF | Granulocyte-macrophage colony-stimulating factor | 0.045 | ns |
| Cytokine | IL-1α | Interleukin-1 alpha | -0.218 | ns |
| Cytokine | IL-5 | Interleukin-5 | 0.179 | ns |
| Cytokine | IL-7 | Interleukin-7 | 0.108 | ns |
| Cytokine | IL-15 | Interleukin-15 | 0.431 | 0.003 |
| Cytokine | IL-16 | Interleukin-16 | -0.045 | ns |
| Cytokine | IL-17A | Interleukin-17 alpha | 0.047 | ns |
| Chemokine | Eotaxin | Eotaxin-1 (CCL11) | 0.089 | ns |
| Chemokine | MIP-1β | Macrophage Inflammatory Proteins (MIP)-beta | -0.147 | ns |
| Chemokine | Eotaxin-3/CCL-26 | Chemokine (C-C motif) ligand 26 | 0.250 | ns |
| Chemokine | TARC | Thymus and activation-regulated chemokine | 0.181 | ns |
| Chemokine | IP-10 | Interferon gamma inducible protein-10 | 0.044 | ns |
| Chemokine | MIP-1α | Macrophage Inflammatory Proteins (MIP)-alpha | 0.242 | ns |
| Chemokine | IL-8 | Interleukin-8 | 0.027 | ns |
| Chemokine | MCP-1 | Monocyte Chemoattractant Protein-1 | 0.098 | ns |
| Chemokine | MDC | Serum Macrophage-derived Chemokine | 0.030 | ns |
| Chemokine | MCP-4 | Monocyte Chemoattractant Protein-1 | 0.158 | ns |
| Angiogenesis | VEGF | Vascular Endothelial Growth Factor | -0.086 | ns |
| Angiogenesis | VEGF-C | Vascular Endothelial Growth Factor-C | 0.248 | ns |
| Angiogenesis | VEGF-D | Vascular Endothelial Growth Factor-D | -0.086 | ns |
| Angiogenesis | Tie-2 | The tyrosine kinase receptor TIE-2 | -0.032 | ns |
| Angiogenesis | Flt-1 | FMS-Like Tyrosine kinase 1 | 0.141 | ns |
| Angiogenesis | bFGF | Basic Fibroblast Growth Factor | -0.052 | ns |
| Proinflammatory | IFN-γ | Interferon-gamma | 0.013 | ns |
| Proinflammatory | IL-1β | Interleukin-1-beta | -0.013 | ns |
| Proinflammatory | IL-4 | Interleukin-4 | 0.134 | ns |
| Proinflammatory | IL-6 | Interleukin-6 | 0.133 | ns |
| Proinflammatory | IL-10 | Interleukin-10 | -0.038 | ns |
| Proinflammatory | IL-12p70 | Interleukin-12p70 | 0.378 | 0.010 |
| Proinflammatory | IL-13 | Interleukin-13 | 0.148 | ns |
| Proinflammatory | TNF-α | Tumour Necrosis Factor-alpha | -0.103 | ns |
| Vascular inflammation | SAA | Human Serum Amyloid A | 0.128 | ns |
| Vascular inflammation | C-reactive protein | C-reactive protein | 0.203 | ns |
| Vascular inflammation | VCAM-1 | Vascular Cell Adhesion Molecule-1 | 0.075 | ns |
| Vascular inflammation | ICAM-1 | Intercellular Adhesion Molecule-1 | 0.144 | ns |

* The correlations were assessed using non-parametric Spearman test (two-tailed).

**Supplementary Table 5. Characteristics of ART-treated participants. Samples from these participants were used for *in vitro* co-culture assays.**

|  |  | PLWH on ART |
| --- | --- | --- |
| Group size | n | 7 |
| Risk factors and demographics | Age (years),  median (range) | 52 (34-59) |
|  | Sex (n)  female: male | 2:5 |
| HIV-1 parameters | Log_10_ HIV-1 VL, median (range) | <50 |
|  | CD4^+^ T-cell count, median (range) | 717 (430-940) |
|  | ART treatment | On ART with undetectable VL |

**Supplementary Table 6. Clinical characteristics of future controllers and non-controllers.**

|  |  |  | Future controllers | Non-controllers |
| --- | --- | --- | --- | --- |
| Group size |  |  | 7 | 20 |
| HIV-1 subtype | Viral subtype |  | A | A |
| Days post estimated date of infection (EDI) | 1 month | Days post-EDI, median (range) | 30 (28–41) | 31.5 (19–42) |
| Days post estimated date of infection (EDI) | 3 months | Days post-EDI, median (range) | 90 (84–100) | 86.0 (66–110) |
| Recruitment clinical site | Research centre, country | Kigali, Rwanda, n (%) | 4 (57.1) | 6 (30.0) |
|  |  | Kilifi, Kenya, n (%) | 3 (42.9) | 14 (70.0) |
| Risk factors and demographics | Age | Age (years),  median (range) | 29 (19–48) | 25 (19–52) |
|  | Sex | Sex (n)  female:male | 1:6 | 2:18 |
|  | Risk group | Discordant couple (DC), n (%) | 4 (57.1) | 6 (30.0) |
|  |  | men who have sex with men (MSM), n (%) | 3 (42.9) | 12 (60.0) |
|  |  | Other heterosexual,  n (%) | 0 (0.0) | 2 (10.0) |
| HIV-1 parameters | 1 month | Log_10_ HIV-1 VL,  median (range) | 4.22 (2.79-5.60) | 5.06 (3.54-7.12) |
|  |  | CD4^+^ T-cell count,  median (range) | 599 (409-1759) | 518 (189-915) |
|  | 3 months | Log_10_ HIV-1 VL,  median (range) | 3.91 (2.79-4.99) | 4.89 (3.64-6.03) |
|  |  | CD4^+^ T-cell count,  median (range) | 637 (371-2000) | 420 (267-1776) |
|  | Set-point VL | Log_10_ pVL set-point,  median (range) | 3.76 (1.69-4.59) | 4.83 (1.83-5.70) |
| HLA-B variants | HLA-B*57 | n (%) | 1 (14.3) | 0 (0.0) |
| HLA-B variants | HLA-B*14 | n (%) | 2 (28.6) | 1 (5.0) |
| HLA-B variants | HLA-B*27 | n (%) | 0 (0.0) | 1 (5.0) |
| HLA-B variants | HLA-B*52 | n (%) | 0 (0.0) | 0 (0.0) |
| HLA-B variants | HLA-B*81 | n (%) | 0 (0.0) | 0 (0.0) |
| HLA-B variants | HLA-B*44 | n (%) | 0 (0.0) | 0 (0.0) |
| HLA-B variants | Any protective HLA-B allele ^*^ | n (%) | 3 (42.8) | 2 (10.0) |

** Any protective HLA-B allele: The combined protective HLA-B allele row includes HLA-B*57, HLA-B*14, HLA-B*27, HLA-B*52, HLA-B*81, HLA-B*44.*

** Twenty non-controllers were included in the analysis; however, one of the non-controllers had NK cell data at three months but not at one month. Therefore, the analysis included 19 non-controllers at one month and 20 at three months.*

**Supplementary Table 7. Antibodies used in flow cytometric analysis.**

| **Antigen/mAb** | **Fluorochrome** | **Supplier** | **Identifier** | **Clone** | **Dilution** | **Staining** |
| --- | --- | --- | --- | --- | --- | --- |
| CD107a | APC/H-7 | BD Biosciences | Cat # 561343 | Clone # H4A3 | 1 in 200μl | During stimulation |
| CD14 | BV510 | BioLegend | Cat # 301842 | Clone # M5E2 | 0.25 in 50μl | Surface |
| CD158b1/b2 | APC | Beckman Coulter | Cat # A22333 | Clone # GL183 | 1 in 50μl | Surface |
| CD16 | BB700 | BD Biosciences | Cat # 746199 | Clone # 3G8 | 0.25 in 50μl | Surface |
| CD2 | BV711 | BioLegend | Cat # 300231 | Clone # RPA-2.10 | 0.5 in 50μl | Surface |
| CD25 | PE-Cy-7 | BD Biosciences | Cat # 335824 | Clone # 2A3 | 1 in 50μl | Surface |
| CD3 | BV605 | BD Biosciences | Cat # 317322 | Clone # OKT3 | 0.5 in 50μl | Surface |
| CD3 | PerCP-eFluor™ 710 | eBioscience | Cat # 46-0037-42 | Clone # OKT3 | 0.5 in 50μl | Surface |
| CD38 | BV785 | BioLegend | Cat # 303530 | Clone # HIT2 | 1 in 50μl | Surface |
| CD4 | BV510 | BioLegend | Cat # 300546 | Clone # RPA-T4 | 0.25 in 50μl | Surface |
| CD56 | BV605 | BD Biosciences | Cat # 562780 | Clone # NCAM16.2 | 1 in 50μl | Surface |
| CD57 | BV421 | BD Biosciences | Cat # 563896 | Clone # NK-1 | 0.25 in 50μl | Surface |
| CD57 | PE/Dazzle 594 | BioLegend | Cat # 359620 | Clone # HNK-1 | 0.25 in 50μl | Surface |
| CD7 | PE/Cyanine5 | BioLegend | Cat # 343110 | Clone # CD7-6B7 | 0.25 in 50μl | Surface |
| FcεRI | FITC | Milli-Mark | Cat # FCABS400F | Polyclonal | 0.25 in 50μl | Intracellular |
| Granzyme B | Alexa Fluor®700 | BD Biosciences | Cat # 560213 | Clone # GB11 | 0.5 in 50μl | Intracellular |
| IFN-γ | BV421 | BD Biosciences | Cat # 562988 | Clone # B27 | 1 in 50μl | Intracellular |
| Ki-67 | FITC | BD Biosciences | Cat # 300231 | Clone # B56 | 3 in 50μl | Intracellular |
| KIR2DL1/KIR2DS5 | APC | R&D systems | Cat # FAB1844A-100 | Clone # 143211 | 1 in 50μl | Surface |
| KIR3DL1/ CD158e | APC | Miltenyi Biotec | Cat # 130-092-474 | Clone # DX9 | 1 in 50μl | Surface |
| KIR3DL2/ CD158K | APC | R&D systems | Cat # FAB2878A-100 | Clone # 539304 | 1 in 50μl | Surface |
| Live Dead stain | Aqua | Life technologies | NA | NA | 1 in 800μl | Surface |
| NKG2A/CD159a | PE-Cy-7 | Beckman Coulter | Cat # B10246 | Clone # Z199 | 0.5 in 50μl | Surface |
| NKG2C/CD159c | PE | R&D systems | Cat # FAB138P-100 | Clone # 134591 | 5 in 50μl | Surface |
| PLZF | PE-CF594 | BD Biosciences | Cat # 565738 | Clone # R17-809 | 0.5 in 50μl | Intracellular |
| Siglec-7 | APC/Fire-750 | BioLegend | Cat # 339207 | Clone # 434 | 0.5 in 50μl | Surface |
| TIGIT | PE | eBioscience | Cat # 12-9500-42 | Clone # MBSA43 | 0.5 in 50μl | Surface |
| TNF-α | BV711 | BioLegend | Cat # 502940 | Clone # MAb11 | 1 in 50μl | Intracellular |
